# Neural Encoding through Hierarchical Amplitude Modulation

**DOI:** 10.1101/2025.11.03.686310

**Authors:** Giulio Ruffini

## Abstract

Hierarchical encoding is a structural element of the Free Energy Principle and related information-centric accounts of brain function, but a concrete circuit-level mechanism for it remains elusive. Here we examine Hierarchical Amplitude Modulation (HAM). In this computationally grounded scheme, information is encoded in the envelope of a carrier and slower brain rhythms multiplicatively modulate the amplitude of faster rhythms, creating a cascade of nested oscillatory envelopes. This multiplicative architecture naturally produces intermodulation frequencies (of the form *f*_*c*_ + Σ*a*_*i*_*f*_*i*_). It predicts that oscillation frequency bands should be log-spaced (*r* ≳ 2–3, depending on the number of modulation layers) to avoid spectral overlap, as in constant-Q filter banks, consistent with the observed logarithmic spacing of canonical brain rhythms (with ratios ∼2–3). HAM’s log-uniform distribution yields a 1*/f* global spectral profile and 1*/f* ^*α*^ sideband spectra in the broadband “aperiodic” component, with the slope determined by modulation depth and band ratio. We then demonstrate modulation and demodulation in the laminar neural mass model (LaNMM), where a fast excitatory-inhibitory oscillator circuit couples with a slower cortical oscillator (Janse-Rit). Through the network’s intrinsic nonlinearities and cross-frequency coupling, amplitude modulation and demodulation are implemented. These results provide a novel circuit-level mechanism for hierarchical predictive coding, linking theoretical principles to observed spectral features of brain activity.

**Highlights:**

- We introduce HAM^1^: a principle for the organization of hierarchical information processing in the brain. Information is encoded through amplitude modulation, where faster rhythms are multiplicatively modulated by slower processes. Information can thus live in signals or their envelopes (and envelopes of envelopes).
- The required intermodulation structure for band protection, *f*_*c*_ ± Σ*a*_*i*_*f*_*i*_, finds a natural implementation in near geometric (log) spacing with a frequency ratio *r* ≳ 2–3 (strong super-increasing hierarchy), yielding constant fractional bandwidths and enabling staged demodulation.
- In addition to geometric frequency spacing, HAM provides two routes to 1*/f* ^*α*^: (i) a cascade link *α* = 2 ln(2*/m*)*/* ln *r*; (ii) a log-uniform mixture that yields 1*/f* in expectation with constant-Q kernels.
- We provide a proof of concept modulation and demodulation implementation in NMM using the LaNMM with PING-like fast generators as carriers and JR-like slow generators for envelope extraction.

## 1 Introduction

Predictive coding has emerged as a central framework for understanding neural processing, positing that the brain continuously generates predictions about incoming sensory data and updates these predictions by minimizing prediction errors.^1, 2^ In the Free Energy Principle, this hierarchical exchange of top-down predictions and bottom-up errors minimizes free energy.^3, 4^ Kolmogorov Theory (KT) embeds this process within Algorithmic Information Theory, suggesting that both biological brains and digital agents optimize predictions by constructing compressive models of the world.^5–10^ All such information-centric proposals of brain function fundamentally rely on encoding information in brain signals to compute prediction errors and update internal representations.

Neural coding schemes have long been debated, with proposals ranging from rate coding (where information is carried by average firing rates) to various temporal strategies—including spike timing, synchrony, phase coding, and burst coding—to capture the richness of neural information.^11–13^ In this context, it is increasingly clear that information is encoded not only in the raw oscillatory signals but also in their amplitude envelopes.^14^ This dual coding strategy is reminiscent of amplitude modulation (AM) in radio communications, where a high-frequency carrier encodes information via its modulated envelope.^15^

In parallel, two empirical regularities stand out in neural field recordings: an aperiodic 1*/f* ^*α*^ background and a near-logarithmic spacing of canonical oscillation bands. Both patterns recur across species and recording modalities, and the aperiodic component can be cleanly separated from periodic peaks in modern parameterizations.^16–21^

Oscillatory activity in the brain naturally segregates into functional bands—such as delta, theta, alpha, beta, and gamma. These canonical ‘bands’ should be understood as families of preferred modes. Although not universally sharp peaks in raw Power Spectra Densities (PSDs), their separability strengthens after accounting for the 1*/f* background or analyzing burst-wise dynamics, and their centers align with an geometric progression across circuits and species. They exhibit a geometric progression with a common ratio *r* ≈ 2–3 (equivalently, equal spacing on a log-frequency axis; constant-Q filterbank,^18^ see Figure 1). That is, the center frequencies increase by a constant multiplier, or equivalently, they are evenly spaced when plotted on a logarithmic scale.^19, 22^ The reason for this spacing is unclear. A leading account is that the approximately logarithmic (geometric) spacing emerges because a constant ratio between bands lets excitability windows and integration times scale with the oscillation period, enabling circuits with heterogeneous axonal/synaptic delays and sizes to coordinate; in this view, the “family” of bands helps overcome delay-imposed processing limits.^19^ Complementarily, it has been proposed that placing bands at equal distances on a log-frequency axis minimizes mutual entrainment/cross-talk and reflects an underlying hierarchical network architecture.^23^

It is believed that oscillatory activity plays a key role in organizing and transmitting information. Gamma oscillations (30–100 Hz) have been implicated in local information processing, feature binding, and communication through coherence,^24–27^ while slower rhythms (theta 4–8 Hz and alpha 8–12 Hz) are associated with attentional gating and top-down signaling.^28–30^

**Figure 1.1:**
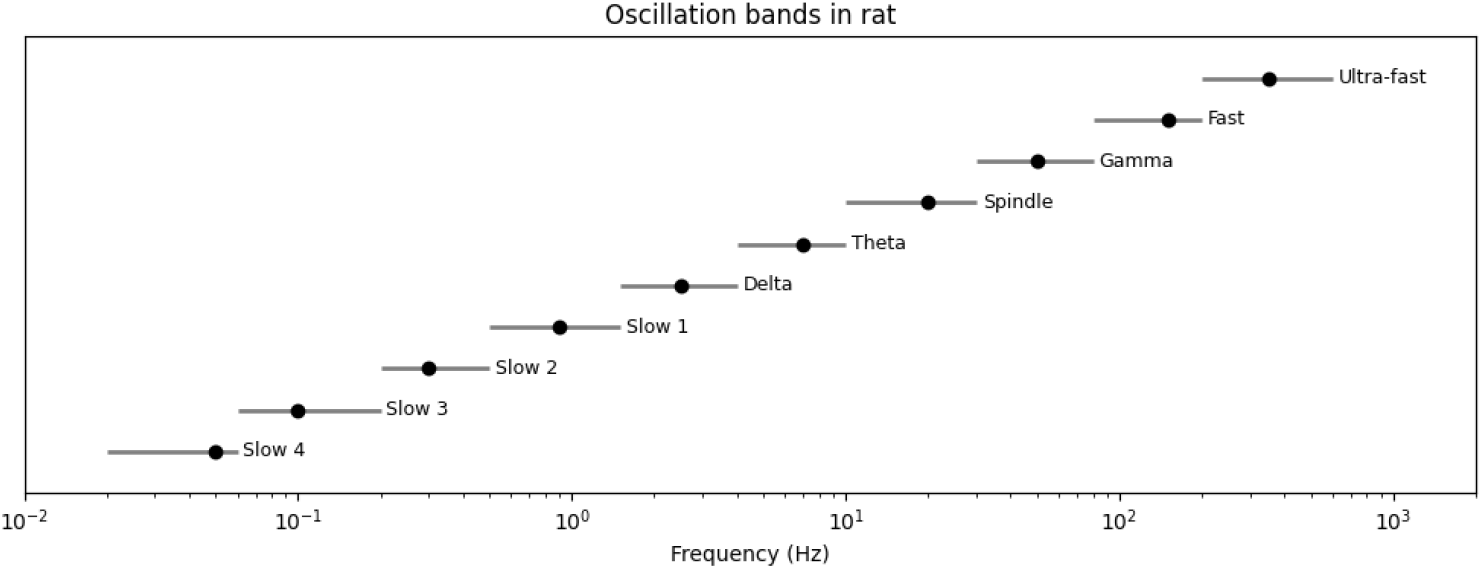
Logarithmic oscillatory hierarchy. Oscillation bands are approximately log-spaced with an approximately common ratio *r* so that *f*_*k*+1_ = *f*_*k*_*/r* (empirically *r* ≈ 2–3 between neighboring bands).^19^ This approaches the theoretical limit of *r* ≳ 3 for non-overlapping side-band clusters and is equivalent to equal spacing on a log-frequency axis (corresponding to a constant-Q arrangement).

Slower oscillations, with longer periods, provide a broad temporal window and allow integration over larger spatial extents with more variable synaptic delays, whereas faster oscillations offer more precise, spatially limited representations of information. Such evidence reinforces the notion that brain oscillations are organized into a conserved, hierarchical set of timescales, with adjacent rhythm classes separated by roughly constant frequency ratios on the order of 2 to 3.^19, 19, 20, 22, 31, 32^

In line with hierarchical processing concepts, such findings suggest that neural information processing is inherently multiscale, with nested oscillatory hierarchies that multiplex information much like modern communication systems. Gamma oscillations, in particular, serve as carriers of sensory information, while slower rhythms modulate these carriers via cross-frequency coupling. Motivated by this, we propose a *hierarchical amplitude modulation* (HAM) framework to describe neural information encoding where information is represented in a layered fashion: the raw oscillatory signal (the carrier), its amplitude envelope, and, recursively, the envelope of that envelope. This nested modulation approach yields a coarse-grained, multiscale, natural representation of information consistent with hierarchical predictive coding. As each successive envelope captures fluctuations at a slower timescale, the system exhibits a logarithmic frequency spacing and power law of oscillatory dynamics—phenomena that align with the observed geometric progression of brain rhythms.

In the following sections, we present the mathematical formulation of the model and demonstrate how it can account for the observed logarithmic spacing and provide a mechanism for the power-law distribution of oscillatory bands. To demonstrate the plausibility of this framework, we then provide a proof-of-concept implementation of amplitude modulation in the Laminar Neural Mass Model (LaNMM),^14, 33–35^ a dual frequency model capturing the interaction between superficial fast and deeper slow oscillations through mechanisms such as phase-amplitude coupling and amplitude-amplitude anticorrelation.

**Figure 2.1:**
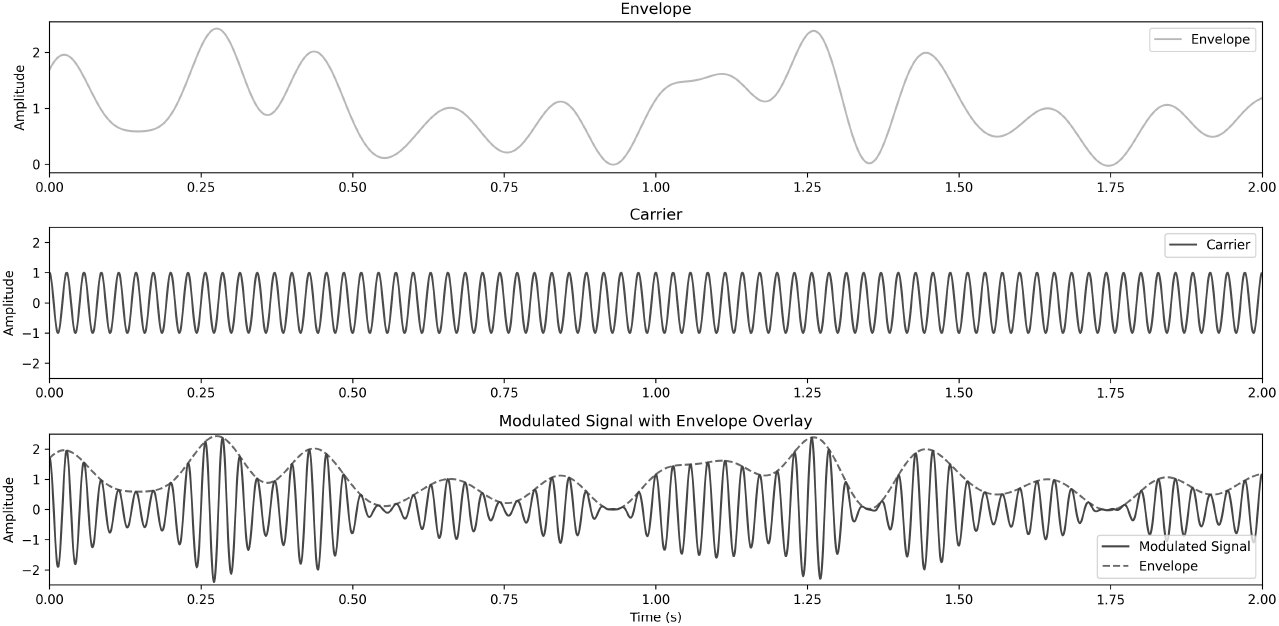
Amplitude modulation (AM), where information in a slower signal is encoded in the envelope of a faster carrier, as in AM radio.

## 2 Hierarchical amplitude modulation (HAM)

### 2.1 Construction

In standard amplitude modulation (AM) (as used in radio), information is encoded in the *amplitude* of a high-frequency carrier wave. For example, an audio signal *m*(*t*) (low-frequency information) can modulate a higher-frequency carrier *C*(*t*) = *A*_*c*_ cos(2*πf*_*c*_*t*) by varying the carrier’s amplitude in proportion to *m*(*t*). The result is a modulated signal

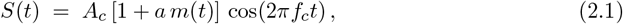

where *a* is the modulation index (chosen so that 1 + *a m*(*t*) > 0 to avoid distortion). Demodulation of *S*(*t*) recovers the original message *m*(*t*) by detecting the envelope. In neural terms, one can think of fast oscillations (e.g., gamma rhythms) acting as carriers whose amplitudes are controlled by slower processes (e.g., theta or delta rhythms).

While classical AM involves a single modulating signal (Fig. 2.1), the brain exhibits *nested* oscillations: faster rhythms modulated by slower rhythms, which in turn are modulated by even slower fluctuations. *Hierarchical Amplitude Modulation* (HAM) generalizes the AM concept to multiple layers of amplitude modulation across different frequency bands. In a HAM cascade, a high-frequency oscillation’s amplitude is modulated by a slower oscillation; this slower oscillation’s amplitude can itself be modulated by an even slower process; and so on, creating a multilevel envelope hierarchy. In effect, information can be carried not only by the fast “carrier” signal, but also by its envelope, the envelope-of-envelope, etc., forming a multiscale encoding of information. Not all frequencies need to be involved in a cascade, but all such combinations should be possible.

#### Nested modulation across oscillatory bands

Consider a chain of *N* nested modulators with descending center frequencies *f*_0_ *> f*_1_ *>* · · · *> f*_*N*_ . For concreteness, let *f*_0_ be a high-frequency carrier and *f*_1_, *f*_2_, … be progressively slower oscillatory components. A depth-*N* HAM signal can be written as a product of an *N* -layer envelope and the carrier,

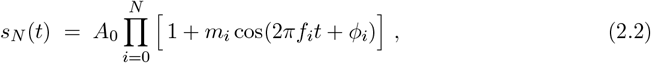

with 0 *< m*_*i*_ *<* 1, the modulation depths at each layer. Here, *i* = 0 corresponds to the fastest oscillation (the putative “carrier”), and each *i >* 0 term represents an oscillatory envelope imposed on the previous layers. All the terms in the product are positive to reflect the nature of firing rates, but this is not a crucial element. Expanding the product in Eq. (2.2) and using the trigonometric identity 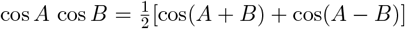 iteratively yields a superposition of cosines at various frequencies, revealing a rich *intermodulation spectrum*: new frequency components (sidebands) appear at linear combinations of the base frequencies. In fact, the spectrum of *s*_*N*_ (*t*) contains sinusoidal components at frequencies

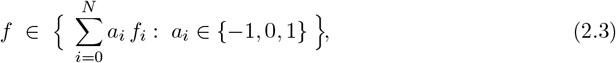

i.e. at the carrier frequency ±*f*_0_, the modulation frequencies ±*f*_1_, ±*f*_2_, …, and all combinations such as *f*_0_ ± *f*_1_, *f*_0_ ± *f*_2_, *f*_0_ ± *f*_1_ ± *f*_2_, etc. Each modulation layer thus spreads the signal’s spectral energy into new sideband peaks spaced around the original frequencies. Importantly, the strength of higher-order sidebands (involving multiple *m*_*i*_ factors) is suppressed for small *m*_*i*_; thus, the cascade remains dominated by its fundamental bands and nearest sidebands if *m*_*i*_ ≪ 1.

This multiplicative, multiband structure is illustrated in Figure 2.2. A high-frequency carrier (Fig. 2.2, top) is successively modulated by a slower oscillation, then by an even slower one, and so on, producing a final complex signal with amplitude fluctuations at multiple timescales (middle). The corresponding frequency spectrum (Fig. 2.2, bottom) displays the original carrier line and a comb of sidebands generated by each modulation layer. Notably, the spectral power decays at lower frequencies, reflecting how each envelope layer contributes a factor *m*_*i*_ *<* 1 to the amplitude (as discussed below). The nested HAM signal can be demodulated in stages to recover the slower envelope signals from the faster carrier, analogous to how a radio receiver detects a single AM envelope – but here performed iteratively across scales.

### 2.2 Avoiding spectral overlap

A critical challenge in any multi-layer modulation is avoiding spectral *overlap* between components. If the newly generated sideband frequencies crowd into neighboring frequency bands, information at different layers would interfere, and demodulation would not be possible. In engineered communication systems, this is solved by allocating *guard bands* and using a *constant-Q* spacing of carriers – meaning each carrier’s bandwidth is a fixed fraction of its center frequency (a logarithmic progression of center frequencies). Strikingly, brain oscillations also appear roughly log-spaced in frequency (each band about a constant ratio higher than the next).^19, 22^ HAM provides a mechanistic reason for this: *geometric spacing* of frequencies naturally prevents intermodulation overlap.

**Figure 2.2:**
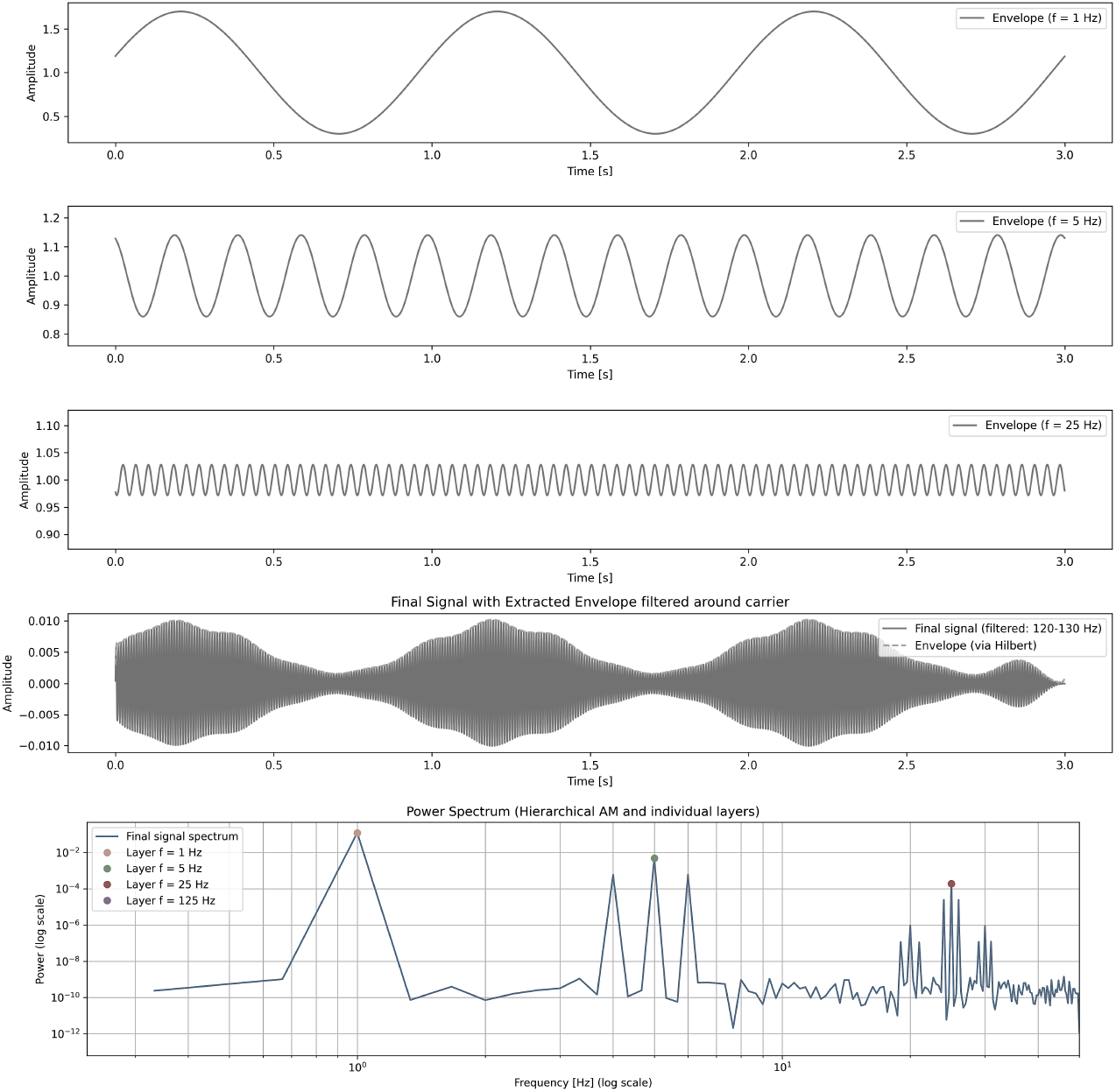
HAM example. A high-frequency carrier is successively modulated by slower oscillations in a nested multiplicative structure. Each subplot presents a key stage in the formation of the hierarchical signal: (A) Envelope Modulations (First three subplots from top): Each line represents an envelope function of the form 1 + *m*_*i*_ cos(2*πf*_*i*_*t* + *ϕ*_*i*_), where each modulation frequency *f*_*i*_ is progressively lower. These envelopes modulate all previously applied layers, creating an intricate structure of amplitude variations at multiple timescales. (B) Demeaned, final Hierarchical AM Signal (fourth plot): The resulting signal after applying all modulation layers. This waveform is the product of all envelope layers modulating a carrier, generating a complex multi-scale amplitude variation. The signal is filtered around the highest carrier frequency (125 Hz). This highlights how the original high-frequency carrier retains structured amplitude variations due to the hierarchical modulation. (C) Spectrum of the full signal. The hierarchical nature of modulation creates a combinatorial spectral expansion, where all integer combinations of modulating frequencies contribute to the final spectrum. This structure requires logarithmic spacing of modulation frequencies to prevent spectral overlap and ensure clear separation of modulation layers. The figure also shows a logarithmic decay in sideband power, which stems from the successive product of *m*_*i*_ factors in each band.

Mathematically, the necessary and sufficient condition to keep all sideband clusters disjoint (and hence ensure sequential demodulation) is to require that each higher-frequency *f*_*k*_ exceeds twice the sum of all lower frequencies (see Appendix A.1). In other words, if

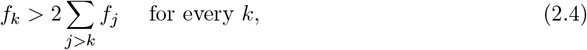

then no combination of slower oscillators can produce a difference-frequency that reaches the *f*_*k*_ cluster. A practical design meeting the necessary condition is to choose a common spacing ratio *r* such that

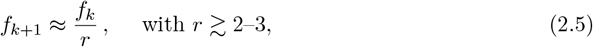

yielding an approximately constant factor separation between successive bands. The minimal ratio needed depends on the number of modulation layers, starting at *r* = 2 for a single AM layer and quickly saturating at *r* = 3 for deeper modulation layers (see Figure A.2 in Appendix A). Geometric spacing (log-uniform in frequency) ensures that each modulation’s sidebands fit into the “gaps” between adjacent bands without overlap if *r* is large enough. Then, the bands behave like well-separated narrow channels. In practice, neural oscillations exhibit *r* ≈ 2–3 (e.g., delta ∼2–4 Hz, theta ∼6–8 Hz, alpha ∼10 Hz, etc.), which provides adequate separation for finite-depth or weak modulation. This log-spaced, constant-Q arrangement mirrors classical wavelet filter banks and the cochlear frequency, and it matches the empirical observation of near-logarithmic spacing in canonical brain rhythm bands.

Crucially, the log-spaced hierarchy means each oscillatory “band” retains its identity (its own frequency range) even as it participates in a larger modulation cascade. By keeping modulation depths *m*_*i*_ moderate (*m*_*i*_ *<* 1), each layer’s sidebands remain as small perturbations around the existing lines. This avoids excessive distortion and allows the original carrier and envelope signals to be separated by bandpass filtering or sequential demodulation. In summary, HAM’s multiplicative encoding across scales naturally favors a logarithmic frequency arrangement: only with such spacing can a deep hierarchy of oscillations coexist without mutual spectral interference.

#### Power-laws

Many electrophysiological signals exhibit an aperiodic 1*/f* -like background in their power spectra. The HAM mechanism offers two complementary explanations for this ubiquitous observation. First, a whole population of log-spaced oscillators will *collectively* produce a 1*/f* -type spectrum in the aggregate. Contributions of equal power per octave (a flat distribution on a log-frequency axis) naturally yield a *P* (*f*) ∝ 1*/f* spectrum. In other words, if oscillatory power is spread roughly evenly across logarithmic frequency bins (as constant-Q band partitioning does), the summed spectrum follows a pink-noise 1*/f* profile (this is a well-known property in audio and wavelet analysis). Second, a single deep cascade of nested oscillations can generate a 1*/f* ^*α*^ spectrum with *α* tuned by neuronal parameters (e.g., modulation strengths and band spacing). We will discuss these two routes next in more detail.

**Figure 2.3:**
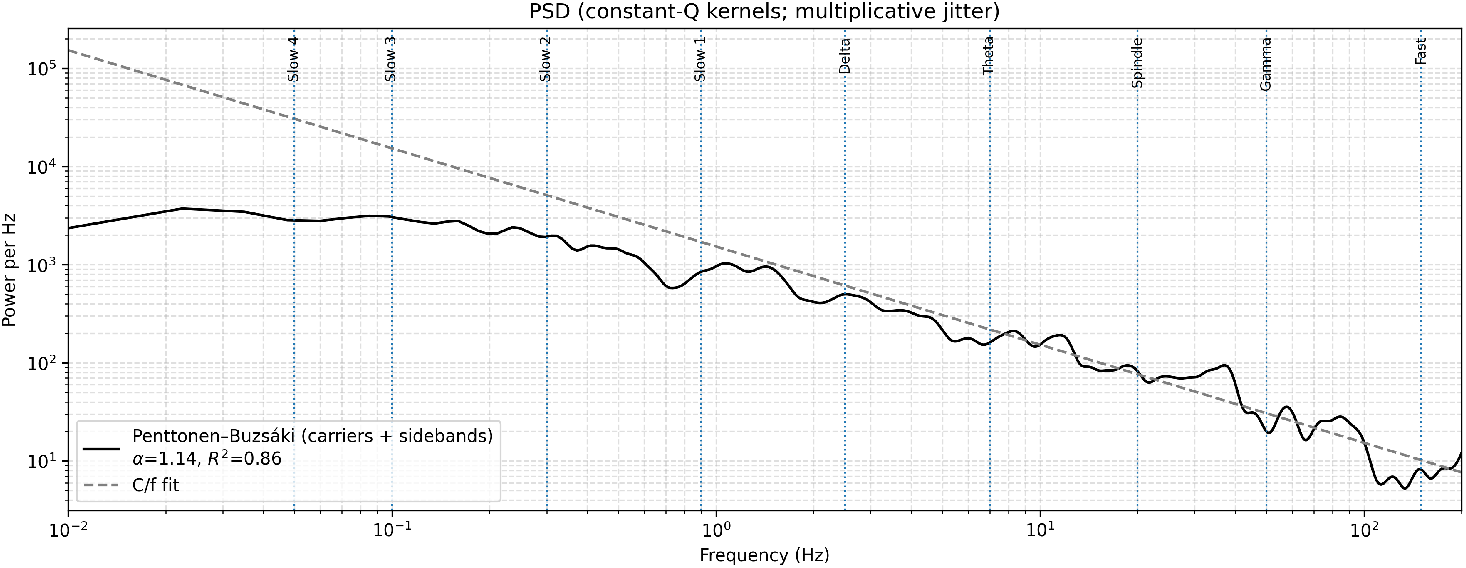
Log–log spectra for log-spaced oscillators with AM sidebands. We display the HAM spectrum using the Penttonen–Buzsáki band centers and equal amplitude carriers, which leads to 1*/f* behavior. The solid curve shows the per-Hz power spectral density (PSD) obtained by summing constant–*Q* Gaussian lines for carriers and their first-order AM sidebands (*f*^*c*^ ± *f*^*m*^, *f*^*m*^ *< f*^*c*^). Each base center is replicated with multiplicative jitter *s* ∼ 𝒰 [1*/*2, 2] to populate each cluster, carrier amplitudes are *A*_*c*_ ≡ 1, and the sideband power is 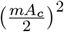 with *m* = 0.9. Dotted vertical lines mark the Penttonen–Buzsáki center frequencies; faint spans indicate their reported ranges. The dashed line is a *C/f* reference. Fitted slopes over 5–250 Hz are indicated in the legend. Because generator density per Hz declines roughly as 1*/f* under log spacing, the aggregate PSD decays with frequency; AM sidebands add a low-*f* pedestal but preserve the 1*/f* backbone in the interior.

#### Global Spectrum

An immediate result from the geometric scaling architecture is power law spectral scaling. As shown in Appendix A.2, a bank of log-spaced oscillators with equal power per band produces *S*(*f*) ≈ 1*/f* . Morevoer, random AM pairings add a constant pedestal; stronger modulation (*m*) or more pairs increase *κ*_0_ and flatten the fitted *α* over finite windows. Finally, using constant-*Q* kernels (or filters) should push the fitted slope in Fig. 1 toward *α* ≈ 1 and increase the fitting range. The assumption *ρ*(*f*) ∝ 1*/f* parallels empirical *logarithmic spacing* of neural bands^19^ and is mathematically equivalent to a constant-*Q* tiling (equal spacing on a log-frequency axis).^36^

We tested the prediction that a log-spaced hierarchy of oscillatory generators produces a per-Hz power density that decays with frequency on a linear axis (reflecting the decreasing generator density per Hz), and that the aggregate spectrum approaches a power law on a log–log axis (see Figure 2.3). We defined a frequency band structure using the set of Penttonen–Buzsaki band centers. To fill gaps on a linear frequency axis, each center *f*_*i*_ was replicated *K* times with multiplicative jitter *u* ∼ Uniform[1, 2], producing micro-generators at *uf*_*i*_ (truncated to [*f*_min_, *f*_max_]). Each micro-generator contributes a narrowband line shaped by a constant-*Q* Gaussian kernel (fractional width *σ/f* = *q*) so that bandwidth scales with center frequency. We set all carrier amplitudes to *a*_*i*_ = 1 (equal power). To emulate amplitude modulation, we drew *M* random pairs (*f*_*m*_, *f*_*c*_) from the micro-generators with *f*_*m*_ *< f*_*c*_ and added first-order sidebands at *f*_*c*_ ± *f*_*m*_, each weighted by (*m/*2)^2^ in power, where *m* ∈ (0, 1) is the modulation depth. Per-Hz power spectra were computed on a dense linear frequency grid and shown on a log–log axis. A 1*/f* reference (*k/f*) was overlaid by matching the median of *S*(*f*) *f* in an interior band. The aperiodic slope *α* was estimated by ordinary least squares on log_10_ *S*(*f*) vs. log_10_ *f* over a predefined range (5–250 Hz). The design tests the claims that (a) constant-*Q* log spacing implies decreasing spectral power density with frequency, and (b) the aggregate spectrum approximates 1*/f* ^*α*^, with sidebands enhancing low-frequency accumulation while preserving the backbone.

**Figure 2.4:**
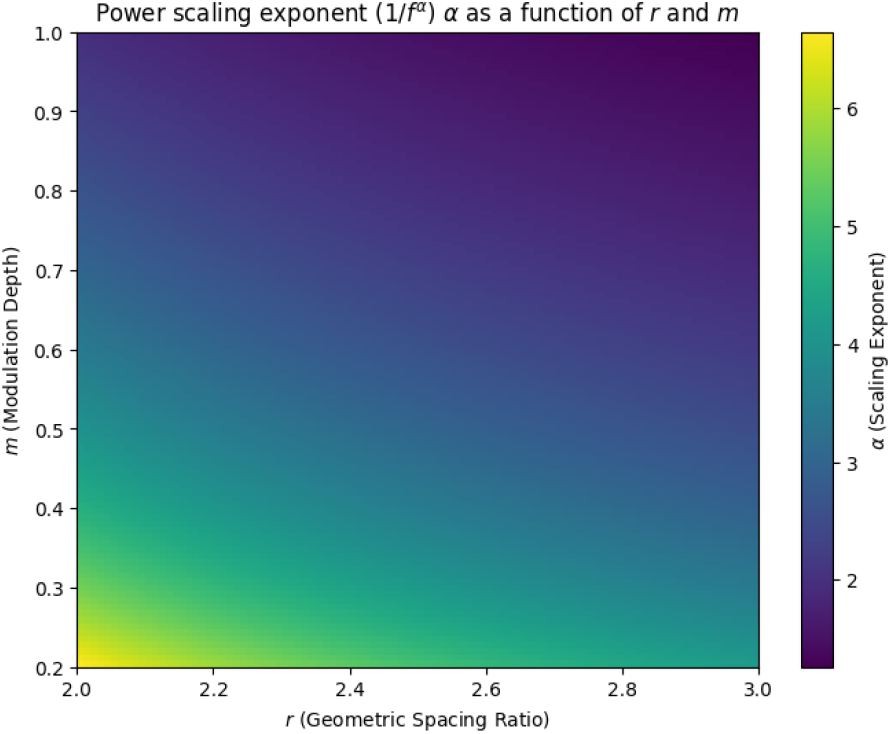
Power-law exponent *α* for log-spaced HAM sideband. For *r >* 1, the toy-model exponent is *α* = 2 ln(2*/m*)*/* ln *r*. Larger *m* (stronger modulation) and larger *r* (wider spacing) both *reduce α* (shallower slope); smaller *m* steepens the slope.

#### Sideband Spectrum

A further consequence of the HAM cascade is the emergence of *power-law* scaling in the sideband spectrum. Because each modulation layer redistributes a fraction of the signal’s power to lower frequencies, the overall spectral density tends to decline with frequency in a scale-free (fractal-like) manner. In fact, under log-spacing, a simple analysis shows that the power spectral density of a hierarchical cascade approximately follows *P* (*f*) ∼ 1*/f* ^*α*^ – a 1*/f* -type law – with the exponent *α* depending on the modulation depth (*m*) and spacing ratio (*r*).

For intuition, consider an idealized case where every layer uses the same depth *m* and ratio *r* for clarity. The first modulation produces sidebands carrying a fraction ∼ (*m/*2)^2^ of the carrier power (since each first-order sideband has amplitude ≈ *m/*2 of the carrier). The next modulation layer acts on those sidebands, generating second-order components of order (*m/*2)^2^ times the first layer’s amplitude, i.e., (*m/*2)^4^ relative to the original carrier’s amplitude, and so on. After *k* layers, the characteristic frequency has decreased to *f*_*k*_ ≈ *f*_0_*/r*^*k*^, and the power at that scale is suppressed by roughly (*m/*2)^2*k*^ relative to the top. Quantitatively, one can show (see Appendix A.2)

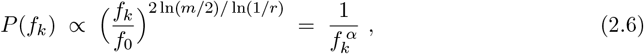

where

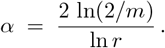

This indicates a 1*/f* ^*α*^ scaling of the cascade’s spectrum. Notably, *α* increases as *m* decreases (weaker modulation yields a steeper spectral drop-off) and *α* decreases as *r* increases (wider band gaps flatten the spectrum). Figure 2.4 illustrates this relationship: stronger modulation (larger *m*) or broader spacing (larger *r*) results in a shallower slope (smaller *α*), whereas small *m* leads to a pronounced 1*/f* ^*α*^ fall-off.

Thus, the emergence of 1*/f* ^*α*^ spectral structure in hierarchical AM systems is explained by two minimal structural constraints: (1) a log-spaced distribution of oscillators and (2) a modulation rule enforcing *f*_*m*_ *< f*_*c*_. Sidebands are not necessary for scale-free behavior but do enhance low-frequency accumulation and steepen the slope.

## 3 Modulation and demodulation in a LaNMM node

We previously introduced a *laminar* neural mass model (LaNMM)^33, 34, 37, 38^ optimized from laminar recordings from the macaque prefrontal cortex^39^ and designed to capture both superficial-layer fast and deeper-layer slow oscillations along with their interactions, which give rise to cross-frequency coupling (CFC).^40^ Here, we show how, thanks to its dual frequency character and nonlinear transfer function, it also provides a candidate circuit implementation of modulation, demodulation, and envelope extraction.

To implement the simplest modulation scheme, two frequency generators are needed. The LaNMM combines conduction physics with two NMMs—Jansen-Rit (P1) and Pyramidal Interneuronal Network Gamma (P2) subpopulations at slow and fast frequencies, respectively—to simulate depth-resolved electrophysiology (see Figure 3.1). Bifurcation properties and cross-frequency coupling have been recently studied.^14, 41^

Modulation and demodulation require a nonlinearity in the model — the transfer function *σ*(·).^38^ This key element in this and related models transduces the sum of membrane potential perturbations *v*(*t*) =Σ^*s*^ *u*_*s*_(*t*) into a firing rate *r*(*t*),

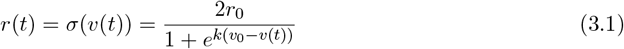

where *u*_*s*_ are the membrane perturbations, *r*_0_ is half the maximum firing rate, *v*_0_ is the potential at *r* = *r*_0_, and *k* determines the sigmoid’s slope. Nonlinear coupling gives rise to *Signal-Envelope Coupling (SEC)*,^39, 42–46^ the coupling of a slow signal to the envelope of a faster signal. In predictive coding, for example,^14^ a slower rhythm (encoding predictions) modulates the envelope (i.e., overall magnitude) of faster oscillations (encoding sensory data), aligning the processing of fast inputs with the slower predictive signal.

Consider such a LaNMM sigmoid node in P1 or P2 acting on the sum of a slow input *s*_1_(*t*) and a narrowband fast input *s*_2_(*t*) ≈ *A*_*c*_ cos(2*πf*_*c*_*t*). The inputs may come from other populations or recurrently from the same population hosting the sigmoid. A local Volterra/Taylor expansion around the operating point gives

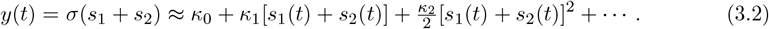

The quadratic cross-term *κ*_2_ *s*_1_(*t*)*s*_2_(*t*) implements amplitude modulation (AM): by the modulation (frequency-shift) property of the Fourier transform, multiplication by cos(2*πf*_*c*_*t*) translates the spectrum of the slow modulator to symmetric sidebands around ±*f*_*c*_ ^47^. Under spectral separation (Bedrosian conditions), the analytic-signal envelope of the band-passed carrier recovers *s*_1_(*t*) up to scale, providing a constructive demodulator.^48, 49^ Such multiplicative mixing yields intermodulation (IM) components at *f*_*a*_ ± *f*_*b*_, 2*f*_*a*_ ± *f*_*b*_, … —the spectral fingerprints of AM/demodulation predicted by HAM—robustly observed across visual paradigms.^50, 51^ Functionally, AM at slow–fast interfaces is consistent with cross-frequency coupling accounts of computation/communication^52, 53^ and with nonlinear mixing known from RF/microwave systems.^54^

We next examine a single LaNMM column to evaluate how it can encode (modulate), demodulate and decode a message via HAM. See Figure 3.1 bottom for the spectra of the P1 and P2 membrane potentials (code for the example is provided at Github).

### Modulation

In the example in Figure 3.1, an external slow oscillatory input in the alpha band (mimicking a “prior” or top-down signal) was injected into the deep-layer pyramidal population P1 while the superficial layer P2 was configured to produce a sustained fast oscillation. The slow input modulates the fast oscillation in P2. In this example, the correlation between external input and P1 was ∼0.52, between P1 and P2 envelope ∼0.98, and between input and P2 envelope ∼0.51.

Modulation can be undone. It has been recently shown that the LaNMM provides a candidate mechanism for the neural function of error evaluation—central to predictive coding—by comparing modulated signals across cortical layers.^14^ This implementation of error evaluation is in fact a demonstration of modulation removal — v. Fig. 7 in Ruffini et al. (2025).^14^ Modulation removal (or error evaluation) is successfully carried out by providing an appropriate prior sign-reversed version of the modulating signal. The incoming slow-band-modulated high-frequency carrier is “unmodulated” by an incoming slow signal with the right signal shape (a sign shift with respect to the high-frequency envelope). Prediction with the right prior demodulates the incoming bottom-up signal, leaving only the carrier.

Second and higher order modulation can be implemented in different ways. One is to again use the modulation approach above, but with a LaNMM model in which the P1 population has a slower natural frequency. P1 can then be driven by an appropriate slower signal. The other is to use envelope-envelope coupling as discussed in Ruffini et al (2025),^14^ where it was used to encode precision in predictive coding.

### Demodulation (envelope extraction)

Finally, it is important to be able to extract the envelope of processed signals (the message). This can be achieved with the architecture described in Figure 3.1 (b). In our implemented example, we find that the carrier envelope can be transferred to the P1 membrane potential with a correlation ∼.45.

**Figure 3.1:**
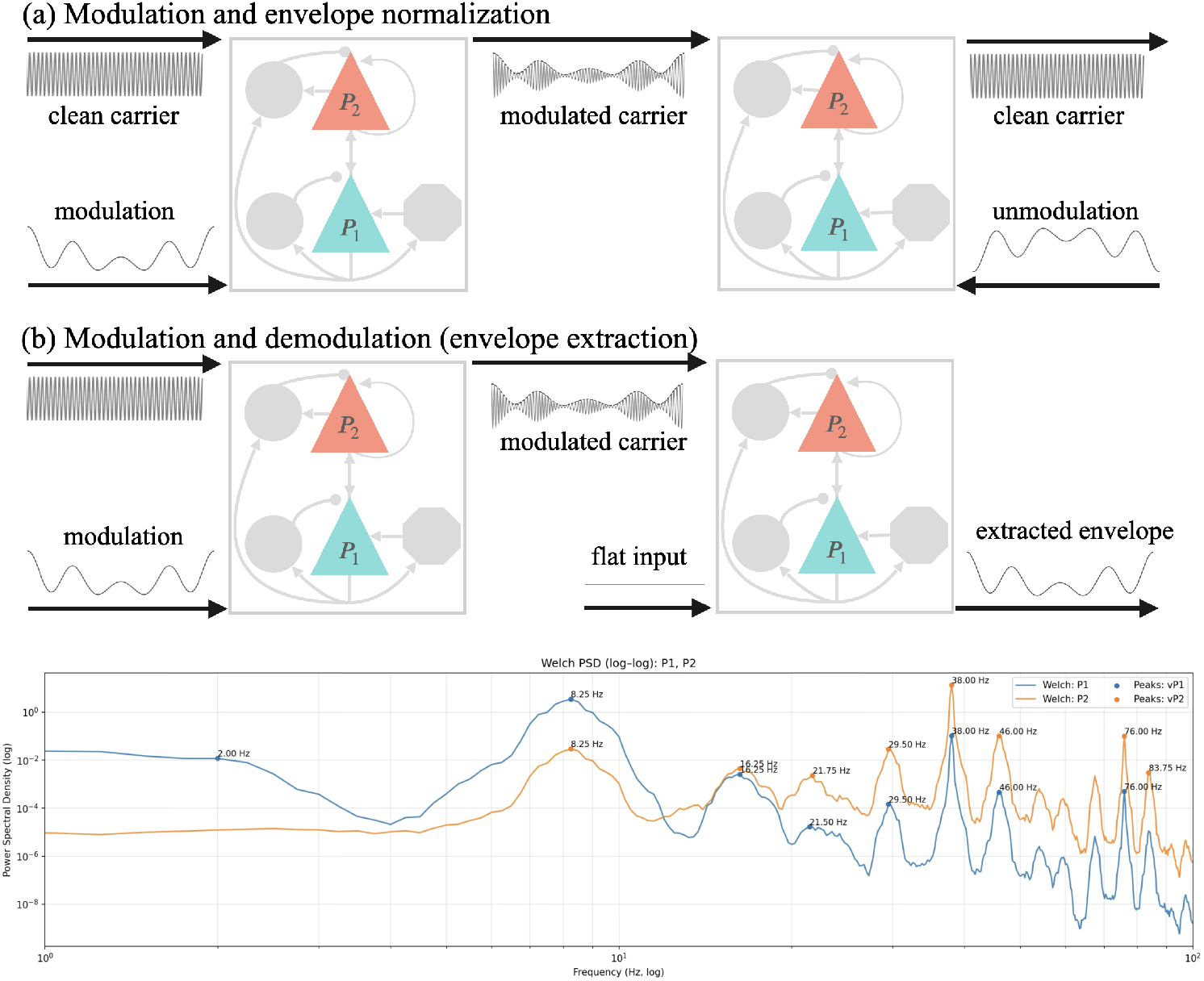
LaNMM architecture for modulation and demodulation. Top (modulation and amplitude normalization): a) The prior modulates the amplitude/frequency of a faster carrier input. The operation is undone to first order by another node with the modulator’s sign-reversed signal (*unmodulation*). b) Modulation and demodulation (envelope extraction). Spectra of LaNMM P1 and P2 (membrane potential) driven by an alpha band input to P1. The alpha-band peak and the sidebands around the gamma peak are clearly visible, as are higher-order terms. **Bottom**: Spectra of P1 and P2 populations during modulation (as in (a)).

## 4 Discussion

### Oscillations in the brain

Oscillations are ubiquitous in neural systems and organize computation across space and time.^20, 22, 55, 56^ Slow rhythms (delta–alpha/beta) tend to coordinate large-scale context and top-down control (feedback), whereas faster beta/gamma rhythms carry local content and bottom-up (feedforward) signals; laminar and inter-areal physiology support this spectral dissociation.^39, 42, 57–59^ In macaque visual cortex, for example, inter-areal Granger-causality in the gamma band is stronger in the feedforward direction (V1 → V4), while alpha-beta influences dominate in the feedback direction (V4 → V1). This aligns with anatomical projections: feedforward pathways originate in superficial layers (rich in gamma) and target layer 4 of upstream areas, whereas feedback pathways arise from deep layers (rich in alpha/beta) and target superficial layers^60^). The result is a frequency-based routing of information, consistent with predictive coding models: fast gamma rhythms convey unexpected input (prediction errors) up the hierarchy, and slower alpha/beta rhythms convey predictions or contextual feedback down.^55^ Cross-frequency coupling (CFC) provides the bridge between these scales, with slow phases modulating fast-band envelopes and timing.^13, 52^

Across EEG/MEG/ECoG and intracranial recordings, canonical rhythm classes (delta, theta, alpha, beta, gamma) occur at progressively higher center frequencies whose separations are roughly constant on a log–frequency axis. In practice, adjacent bands are typically separated by a factor of about two to three (e.g., *θ* ∼ 4–7 Hz, *α* ∼ 8–12 Hz, *β* ∼ 15–30 Hz, *γ* ∼ 40–80 Hz), yielding an approximately geometric hierarchy of timescales.^18, 19, 22, 28, 29, 55^ Penttonen and Buzsáki explicitly quantified near–log spacing in rodents;^19^ convergent human and non-human primate reviews and textbooks report comparable band centers and boundaries, which imply ratios *r* ≈ 2–3 between neighboring classes.^22, 28, 29, 55^ Methodological work that separates periodic peaks from the aperiodic background further confirms that these peaks recur at roughly logarithmic intervals across individuals and tasks.^18^

Recent frequency-tagging work makes this hierarchy directly observable: when two inputs are presented at distinct tagging rates, neural responses exhibit *intermodulation* (IM) components at sums and differences of the tags—clear fingerprints of multiplicative mixing/demodulation predicted by HAM. This has been leveraged in steady-state visual paradigms to probe integration and attention.^61^ Comprehensive reviews document robust IM sidebands across visual cognition ^50^ and characterize their generators and applications ^51^. In our framework, these side-bands arise naturally from envelope–carrier architectural interactions: slow predictive rhythms (alpha/theta) multiplicatively gate fast carriers (beta/gamma), yielding IM structure around each band center while preserving separability under approximate logarithmic spacing (constant-*Q*). This same logic explains why, after constant-*Q* preprocessing, both periodic peaks and an aperiodic 1*/f* ^*α*^ backbone can be read without mutual bias.^17, 18^

### Computation with oscillators

Beyond description, oscillations can *compute*. Analog and hybrid models use coupled oscillators as primitives for signal processing and decision dynamics.^62^ A broader theory and hardware stack for *computing with oscillators* now connects phase, frequency, and amplitude modulation to classification, constraint satisfaction, and optimization ^63^. In AI, recurrent oscillator reservoirs (e.g., Kuramoto-type networks) provide nonlinear fading-memory dynamics and rich readouts for sequence learning and control.^64, 65^ This may provide a route to the large energy savings realized in computational biological systems compared to artificial ones.^66^ Converging theory in systems neuroscience argues that the neocortex itself employs recurrent oscillator networks whose computational capacity depends on the *patterning and heterogeneity* of local oscillators—and that increasing heterogeneity (while avoiding overlap) enhances separability and pushes the network toward powerful operating regimes near criticality.^67^ Large-scale modeling further shows how oscillatory motifs implement routing, working-memory control, and flexible task switching, with function tied to band-specific coupling and multiplexing ^66^.

HAM fits squarely within this computational view: it *requires* oscillator heterogeneity and benefits from log-uniform (constant-*Q*) spacing to prevent IM collisions, enabling layered modulation /demodulation pipelines across the cortex. In practice, this predicts that (i) modest increases in frequency diversity should improve multiplexed readout, and (ii) approximately logarithmic spacing maximizes usable channel capacity—two testable signatures in laminar/ECoG/MEG experiments and neuromorphic oscillator arrays. In HAM, relatively fast rhythms serve as carriers whose amplitudes are multiplicatively structured by slower processes; staged demodulation then makes these envelopes readable downstream. This implements a neural analog of multi-layer AM in communication engineering, but realized in laminar circuits.

### From analog/digital representations to hierarchical envelopes

Information in engineered systems may be encoded digitally (symbolic, discrete) or analogically (continuous waveforms). Neural systems are fundamentally analog at the field level, with information carried both by oscillatory signals themselves and by their amplitude envelopes—the latter being a natural vehicle for slow context that modulates faster content.^22^ Auditory ECoG demonstrates this duality: high-gamma envelopes track the stimulus envelope and support direct speech reconstruction.^68, 69^ HAM formalizes a coarse-to-fine organization in which information can reside in the signal and in successive envelopes (envelope-of-envelope, etc.), yielding a principled multi-scale code. Debates on neural coding (rate, temporal/ spike-timing, phase, and burst codes) converge on the view that multiple encoding schemes coexist and are multiplexed across scales.^11–13^ In this light, HAM specifies *one* concrete way slow rhythms coordinate fast content: theta/alpha phases gate gamma-band carriers (rate/temporal information) via gain control, with nested envelopes enabling compositional, layered messages. This aligns with “communication-through-coherence” and the spectral dissociation of feedforward/feedback pathways.^55, 57, 58^ More broadly, oscillator-based computation is increasingly recognized as a viable computing paradigm; HAM connects that literature to cortical field dynamics.^63, 66, 67^

### A rationale for log spacing

Intermodulation at each stage produces sideband clusters around every center frequency. Avoiding spectral collisions across levels requires the span of slower combinations to fit inside the guard band of faster clusters. A strict super-increasing hierarchy *f*_*k*_ *>* 2Σ _*j>k*_ *f*_*j*_ suffices to keep clusters disjoint; geometric spacing *f*_*k*+1_ = *f*_*k*_*/r* satisfies this with margin for *r* ≳ 2–3 and yields constant fractional bandwidths (constant-*Q*), making demodulation scale-invariant and mirroring classical filterbanks and cochlear tiling.^36^ The minimal ratio depends on the number of modulating layers (*r* = 2 is the minimum for a single carrier/modulator AM arrangement, see Figure A.2 and Appendix A for details). Exact ratios are not essential—a weaker non-overlap inequality already pushes the ensemble toward a log-uniform density of centers. Empirically, *r* ≈ 2–3 lies near the theoretical *r >* 2–3 threshold, explaining why real bands show limited but tolerable overlap.

### Two routes to the aperiodic 1*/f* ^*α*^

HAM predicts 1*/f* ^*α*^ via two complementary mechanisms. (i) *Cascade route:* with log spacing and small, quasi-uniform modulation depth *m, k*-modulator terms scale ∼ (*m/*2)^*k*^ ; because *k* ∼ log_*r*_(*f*_1_*/f*), *P* (*f*) ∝ *f*^−*α*^ with *α* = 2 ln(2*/m*)*/* ln *r*. This links slope to circuit variables, predicting how *α* should shift with changes in *m* or *r*. (ii) *Mixture route:* a log-uniform bank of narrowband (constant-*Q*) oscillators already yields ∼ 1*/f* in expectation (equal power per octave). Random AM pairings preserve this backbone in the interior but induce predictable edge deviations: a small-difference pedestal at low *f* and a finite-support roll-off at high *f* . Analysis choices (edges; constant-Hz vs. constant-*Q* smoothing) determine whether fitted 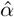 flattens or steepens over finite windows.^17, 18^

### Mechanistic implementation

Here we provided an implementation proof-of-concept of amplitude modulation in the LaNMM, with base frequencies in the alpha and gamma bands, although other frequencies can be implemented in this model^14^ and related ones^70^ by adjusting parameters. In LaNMM, PING-like fast generators and JR-like slow generators map naturally onto “carrier” and “envelope” roles, with demodulation operationalized by analytic-signal envelopes.^49^ These choices are consistent with laminar spectral asymmetries—gamma favoring feedforward content and alpha/beta favoring feedback predictions.^57, 58^ The generation of other frequencies may involve larger-scale mechanisms, e.g., thalamo-cortical loops, but neural mass models have been proposed for most of the canonical bands (see Table A.1.

### Predictions

A core prediction of multiplicative envelopes besides approximate geometric spacing is *intermodulation arithmetic*: sum/difference terms (*f*_1_ ± *f*_2_, 2*f*_1_ ± *f*_2_, …) should emerge when two tagged inputs interact nonlinearly or when demodulation occurs. This is precisely what steady-state visual paradigms report and exploit.^51, 61^ Intermodulation components (IMs) are now used to assay integrative neural mechanisms from low- to high-level vision, attention, and multisensory binding, strengthening the case for HAM-like processing ^50^. In laminar circuits, HAM further predicts superficial carrier-dominant activity coexisting with deeper envelope-dominant readouts, and cross-frequency coupling, matching reported interactions.^57, 58^

HAM makes direct, perturbation-level predictions. (i) *Spacing:* approximately geometric spacing should minimize cluster collisions and facilitate staged demodulation. (ii) *IM scaling:* the amplitude of IM terms scales with products of modulation depths, providing a calibration handle in frequency-tagged designs. (iii) *Laminar dissociation:* superficial layers should carry fast carriers; deeper layers should track their envelopes in prediction-heavy contexts. (iv) *Perturbations:* reducing a slow rhythm’s amplitude should reduce the envelope variance of faster activity it modulates (steepening the PSD), whereas entraining a slow rhythm should increase that variance (flattening the PSD). These can be tested with closed-loop stimulation and preregistered laminar/ECoG/MEG paradigms, coupled to LaNMM-based parameter inference to link controlled changes in *m* and *r* to shifts in *α*. Constant-*Q* preprocessing and explicit separation of periodic vs. aperiodic components are essential to stabilize slope estimates and avoid spurious CFC.^18^

### Relation to alternative accounts

Multiple mechanistic routes can yield the observed near–logarithmic spacing of brain rhythms and their cross-frequency interactions. One line argues that approximately logarithmic spacing follows from hierarchical anatomy and finite conduction delays, whereby integration windows scale with the oscillation period, allowing circuits with heterogeneous axonal/synaptic delays and sizes to coordinate.^71^ In this view, faster rhythms remain local while larger networks synchronize more slowly, consistent with evolutionary preservation of a hierarchy of timescales and size-invariant timing,^32^ and with a posterior–anterior cortical frequency gradient in humans.^72^ Whole-brain models further show that realistic, distance-dependent delays can generate alpha-band phase networks.^73^ Complementarily, spacing as equal steps on a log-frequency axis has been argued to minimize mutual entrainment/cross-talk and to reflect hierarchical network architecture—shown in reverse-engineered networks that reproduce log-spaced spectral peaks.^23^ Recent formal work refines this optimization idea: “golden rhythms” place neighboring centers at the golden ratio to maximize multiplexing (minimal interference) while preserving efficient triadic cross-frequency interaction;^74^ related analyses propose a binary 1:2 hierarchy (and a “golden-mean rule”) linking brain and body oscillations and predicting harmonic/nested coupling across scales. Klimesch’s binary hierarchy theory posits that cross-frequency interactions are governed by two principles—phase–to–amplitude (envelope) modulation and phase–phase (harmonic) coupling—applied to brain and body oscillations arranged on a 1:2 ladder.^31^

A parallel mechanistic route is self-organization: with activity-dependent plasticity, neural populations can split into a few frequency clusters whose slower envelopes modulate faster components, offering a circuit basis for band “quantization”.^75–77^ Methodologically, the apparent discreteness strengthens after modeling out the aperiodic 1*/f* background and analyzing burst-wise dynamics—revealing approximately constant-*Q* spacing of putative peaks^18^—and recent state-space reconstructions report a low-dimensional geometric “core” that organizes band interactions^78^). ‘

Additionally, a large literature treats slow phases modulating fast-band amplitudes (PAC) and related CFC as a control mechanism that routes information and coordinates computations across scales.^52^ Laminar and inter-areal data support a spectral dissociation of feedforward vs. feedback: gamma/theta dominate feedforward, alpha/beta feedback;^57, 79^ large-scale/laminar modeling reproduces these asymmetries,^45^ and a recent laminar atlas shows ubiquitous deep (alpha-beta)/superficial (gamma) motifs across primates.^80^

HAM differs from the above by proposing a compositional information-encoding scheme and by focusing on its consequences for oscillatory spectral structure. It does not derive the spacing primarily from biophysical delays or carrier interference-avoidance; instead, it treats the band hierarchy as a normative, codebook-like quantization that supports clean, scale-invariant multiplexing under explicit constraints, with specific multiplicative schemes and falsifiable predictions about when/where centers can shift while preserving constant-*Q* structure. Our results show how nested envelopes can carry specific information and yield testable spectral consequences. HAM starts from the central role of amplitude modulation but generalizes the spacing: inter-modulation sidebands impose guard-band constraints that are satisfied by *any* approximately geometric (constant-*Q*) spacing with ratio *r* ≳ 2-3. It also offers a constructive path to fractal-like spectra by structurally favoring criticality. It is compatible with signatures of near-critical dynamics (avalanches, scale-invariant correlations). Combining HAM, time-constant, or E–I explanations will require perturbational tests and careful finite-size controls—not spectral fits alone.^81, 82^

Finally, prior work in signal processing has noted that constant-*Q* (log-spaced) filterbanks with equal power per octave imply an overall 1*/f* (“pink-noise”) backbone, and that cascade constructions (e.g., mixtures of relaxation times or multiplicative cascades) naturally generate 1*/f* ^*α*^ spectra.^36, 81, 83–86^ We place that in a neural context and further link it to predictive coding dynamics (with deep layers modulating superficial layers). HAM can be viewed as a mechanistic complement to descriptive models of 1*/f* noise (e.g., fractional Gaussian noise, self-organized criticality). Rather than assuming an ad hoc fractal process, HAM builds 1*/f* from interacting oscillatory components with clear physiological mapping. Beyond qualitative arguments for 1*/f*, HAM derives two mechanistic routes—(i) a multiplicative cascade linking slope *α* to modulation depth *m* and spacing *r*, and (ii) a mixture route from aggregating log-spaced narrowband oscillators.^18^

### Predictive coding link

HAM provides a hierarchical information encoding backbone for predictive coding. As proposed in Ruffini et al. (2025),^14^ in hierarchical predictive coding, deep populations provide slow predictions *p*_*k*_(*t*) while superficial populations carry content in their envelopes. Error evaluation at level *k* is essentially carried out by a signed *demodulation*:

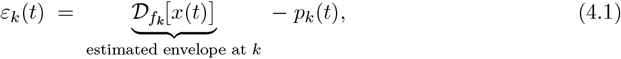

where *x*(*t*) is the bottom-up signal, *p*_*k*_(*t*) the prediction for the *k*-th layer, and D stands for demodulation at frequency *f*_*k*_ — i.e., envelope extraction followed by subtraction (the “Comparator”). Laminar data support this mapping: gamma-band activity dominates feedforward channels and alpha/beta envelopes dominate feedback, consistent with deep-layer envelope readout and staged demodulation across the hierarchy.^57, 58^ Thus, modulation order is immaterial (product structure), whereas unmodulation should proceed fast→slow (or in parallel constant-*Q* channels) to expose the nested envelopes that predictive circuits compare to their top-down forecasts (see Appendix A.1).

### Outlook

Hierarchical Amplitude Modulation offers a coherent explanation linking multiple scales of neural dynamics. By requiring logarithmic frequency spacing and limited modulation depth, HAM ensures that oscillatory bands can nest without losing their identity, providing a multi-layer carrier for information. The framework elegantly accounts for the origin of 1/f spectra as a consequence of either deep cascades or aggregated log-oscillators. It connects to laminar circuit anatomy, suggesting that the cortex’s layered structure is not just a feedforward/feedback arrangement but also a modulation pipeline for neural signals. There are many avenues to build on this work: incorporating more realistic spiking dynamics, exploring how noise and oscillations interplay in HAM (since external noise could either disrupt or be filtered by such a system), and extending to other frequency domains (e.g., could very slow *<* 1 Hz fluctuations modulate high-frequency oscillations across brain-wide networks?). The HAM model provides clear predictions and a language to discuss how the brain’s rhythms might interact to encode information. As neuroscience moves toward understanding the brain as a multi-scale communications network, HAM may provide a valuable piece of the puzzle, bridging theories from engineering with the biological reality of brain oscillations.

## Acknowledgements

The author thanks Francesca Castaldo for helpful discussions and for reviewing the manuscript.

## Code availability

Code for simulations and figures is available at Github.

## A Appendix: Mathematical details

AM radio operates by encoding information in the amplitude of a high-frequency carrier wave. In this context, “information” typically refers to the audio signal we want to transmit, such as music or voice. This information is represented as a low-frequency signal, which modulates or varies the amplitude of the high-frequency carrier wave. The carrier itself oscillates at a frequency much higher than the signal, enabling it to be transmitted over long distances efficiently. During modulation, the carrier’s amplitude is adjusted to follow the waveform of the audio signal, embedding the desired information within the carrier’s envelope. When received, this combined signal undergoes demodulation to extract the original audio information from the carrier. Demodulation detects variations in amplitude and reconstructs the original low-frequency signal. This process highlights how AM radio relies on distinct components: a steady carrier, the modulating information signal, and demodulation to recover the original information for playback. The *carrier signal C*(*t*) can be expressed as *C*(*t*) = *A*_*c*_ cos(2*πf*_*c*_*t*), where *A*_*c*_ is the amplitude of the carrier, and *f*_*c*_ is the carrier frequency. To encode the information, the amplitude of the carrier is varied according to the lower-frequency *information signal m*(*t*), producing the *modulated signal S*(*t*),

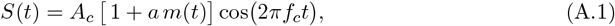

where *a m*(*t*) is scaled such that 1 + *a m*(*t*) remains positive, ensuring no distortion occurs during modulation. Here, *A*_*c*_ is the carrier amplitude, *a* is the *modulation index, m*(*t*) is the baseband (modulating) signal with frequency content up to *f*_*m*_ ≪ *f*_*c*_, and *f*_*c*_ is the carrier frequency. This generic model subsumes both single-tone AM (where *m*(*t*) = cos(2*πf*_*m*_*t*)) and multi-tone/wideband modulations, as long as the maximum modulating frequency is less than *f*_*c*_. At the receiver, a *demodulation* process retrieves the original information signal *m*(*t*) by detecting variations in the amplitude (or envelope) of *S*(*t*).

While standard AM encoding relies on a single layer of modulation, where the information signal *m*(*t*) modulates the amplitude of a high-frequency carrier, this concept can be extended to a *hierarchical framework in which multiple layers of modulation occur at different timescales (HAM*. In HAM, each modulating signal not only modifies the carrier but also modulates the envelope of another modulating signal at a slower timescale, forming a cascade of amplitude modulations across hierarchical frequency bands.

### A.1 HAM and logarithmic spacing of modulating bands

Hierarchical Amplitude Modulation (HAM) refers to a nested modulation scheme in which a high-frequency carrier is amplitude-modulated by a signal, whose amplitude in turn is modulated by another slower signal, and so on across multiple layers. In essence, information is encoded at multiple hierarchical levels of the amplitude envelope. We describe this concept step by step, analyze the resulting spectra at each stage, and show why a logarithmic (geometric) spacing of carrier and modulation frequencies naturally emerges to prevent spectral overlap, as observed in electrophysiological signals^19, 22^ (see Figure 1). Finally, we discuss general insights into how such a structure can be demodulated and generalized.

**Figure A.1:**
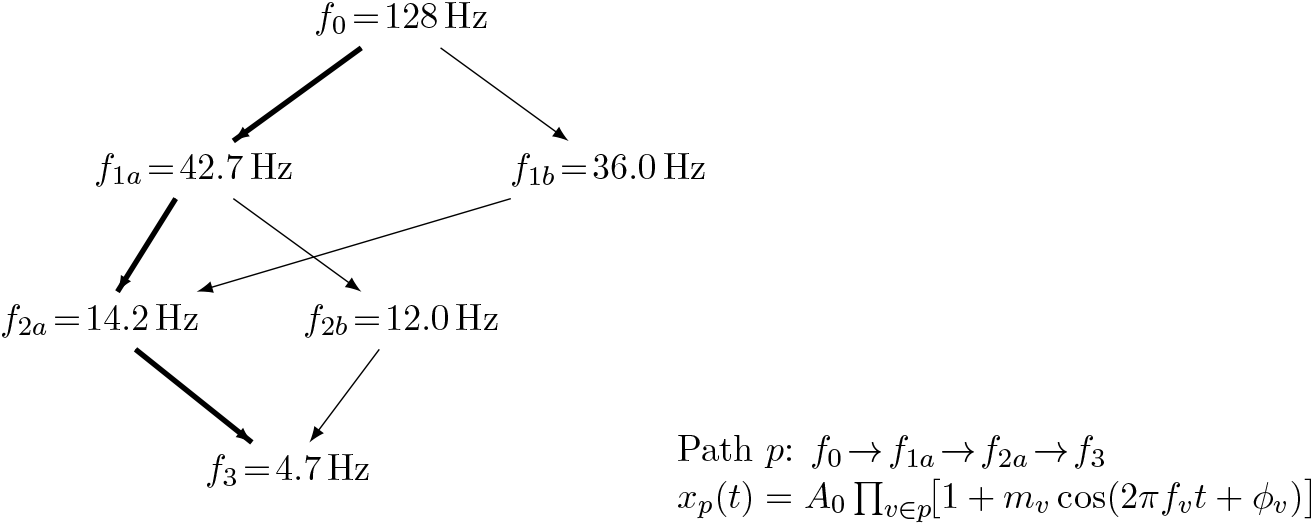
Directed graph of oscillators (nodes) and allowed modulations (arrows). A highlighted path *p* represents one hierarchical AM cascade; the full signal is the sum over all such paths (Eq. (A.6)).

#### Definition (single HAM cascade)

In neural circuits, there is no singled-out “carrier”. We therefore treat the fastest oscillator as any other building block in a multiplicative cascade around a positive baseline. Let the highest frequency (“carrier”) signal be

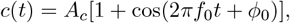

and let *f*_0_ *> f*_1_ *>* · · · *> f*_*N*_ be modulating tones. A *depth-N hierarchical amplitude-modulated signal* is

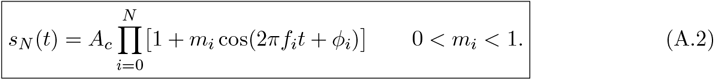

By expanding (A.2) and using the trigonometric identity 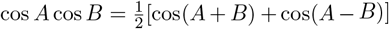, this introduces additional sidebands. The (one-sided) spectrum contains lines at

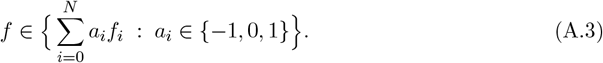

At lowest order (small *m*_*i*_), the first two layers (*i* = 0, 1) contribute ±*f*_0_, ±*f*_1_, and ±(*f*_0_±*f*_1_), and in general, a term involving *k* nonzero coefficients has amplitude proportional to _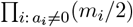_. At depth *N*, the number of algebraic combinations is at most 3^*N*^ ; counting only distinct positive-frequency lines (ignoring DC and mirror symmetry) gives an upper bound ≤ (3^*N*^ −1)*/*2. Degeneracies occur when frequencies are commensurate.

#### Graph-based HAM

Here we analyze the more general case where multiple HAM chains interact. Let *G* = (*V, E*) be a directed acyclic graph (DAG) of oscillators (Fig. A.1). An edge (*v* → *u*) ∈ *E* means that the slower node *v modulates* the faster node *u*. Each node *u* ∈ *V* has a carrier *c*_*u*_(*t*) = cos(2*πf*_*u*_*t* + *ϕ*_*u*_) and gain *A*_*u*_ *>* 0. Multiple incoming modulators *sum first* and then modulate the carrier at *u*.

##### Presynaptic drive and node output

Define the (dimensionless) presynaptic drive

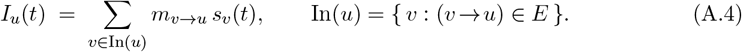

Here *s*_*v*_(*t*) is the signal exported by *v* that serves as a modulator (e.g., a narrowband *s*_*v*_(*t*) = cos(2*πf*_*v*_*t* + *ϕ*_*v*_) or, more generally, the envelope at *v*). The node output is

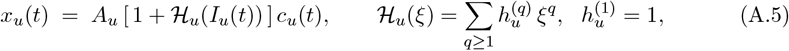

and the network signal is the superposition *S*(*t*) = Σ _*u*∈*V*_ *x*_*u*_(*t*). Using (A.5) and the Taylor series of ℋ _*u*_,

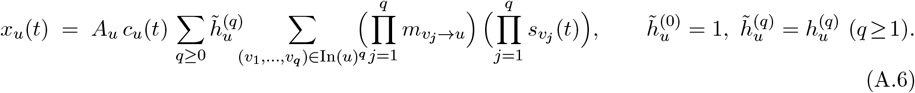

Recursively substituting the same form for each 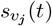 yields a convergent sum over *finite directed forests* in the DAG (finite because *G* is acyclic and *m*_*v*→*u*_ are small), i.e., a global sum of time-domain products whose coefficients are monomials in the edge depths *m*_*e*_.

##### Spectral support

Assume as before that each exported modulator has a narrowband component at *f*_*v*_ (e.g., *s*_*v*_(*t*) = cos(2*πf*_*v*_*t* + *ϕ*_*v*_) at leaves, and more generally *s*_*v*_ inherits the narrowband content of its ancestors). Then every term in (A.6) is a product of cosines. Using 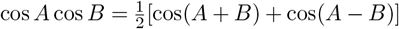 iteratively gives again a spectrum as in Equation A.3. To see this, order the DAG topologically. For each *u* ∈ *V*, let Anc(*u*) be its ancestors. Then all non-DC spectral lines of *S*(*t*) lie in

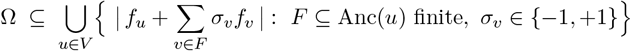

(with mirrored negative frequencies), i.e., *linear combinations with coefficients in* {−1, 0, +1}. That is, Equation A.3 holds. Amplitudes are polynomials in the *m*_*e*_ (products of edge depths on the contributing forests) times node factors 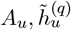.

#### Spacing constraints: necessary and sufficient

Let *f*_1_ *> f*_2_ *>* · · · *> f*_*N*_ *>* 0 and let the sideband cluster around *f*_*k*_ be 𝒞 _*k*_ ⊂ [*f*_*k*_ − *S*_*k*_, *f*_*k*_ + *S*_*k*_] with *S*_*k*_ = Σ _*j>k*_ *f*_*j*_. We now show that {𝒞 _*k*_} are pairwise disjoint iff *f*_*k*_ *>* 2 Σ ^*j>k*^ *f*_*j*_ for all *k*.

##### Theorem A.1. Necessary and sufficient condition for non-overlap of clusters.

*Let f*_1_ *> f*_2_ *>* · · · *> f*_*N*_ *>* 0 *and define the half-width S*_*k*_ := Σ _*j>k*_ *f*_*j*_, *and the (one-sided, positive-frequency) sideband cluster*

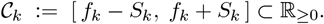

*Then the family* 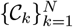*is pairwise disjoint (strictly, i*.*e*., *no touching) if and only if*

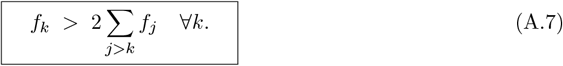

*Proof. Support envelope*. By construction of the HAM spectrum, every line in the “*k*-th cluster” has the form *f* = *f*_*k*_ + Σ _*j>k*_ *a*_*j*_*f*_*j*_ with *a*_*j*_ ∈ {−1, 0, 1}. Hence the smallest/largest possible values in that cluster are obtained by taking all signs negative/positive, and thus

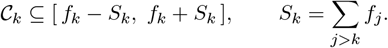

(For the interval proof it suffices to work with these extremal bounds.)

##### Sufficiency

Assume (A.7). Then ∀*k, f*_*k*_ *>* 2*S*_*k*_ and

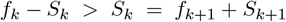

Therefore the entire *k*-th interval lies strictly to the right of the (*k*+1)-st interval: 𝒞_*k*_ ∩𝒞_*k*+1_ = ∅. Since *f*_1_ *>* · · · *> f*_*N*_ orders the intervals on the real line, disjointness of all adjacent pairs implies the whole family {𝒞_*k*_} is pairwise disjoint.

##### Necessity

Conversely, suppose the clusters are pairwise disjoint. In particular, 𝒞_*k*_ and 𝒞_*k*+1_ do not intersect, so

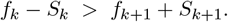

Using *S*_*k*_ = *f*_*k*+1_ + *S*_*k*+1_ this reduces to *f*_*k*_ − *S*_*k*_ *> S*_*k*_, i.e. *f*_*k*_ *>* 2*S*_*k*_ = 2Σ_*j>k*_ *f*_*j*_. Since this holds for each adjacent pair, it holds for all *k*, proving (A.7). □

#### Corollary (Geometric spacing)

If for all *k* (log-uniform or geometric spacing),

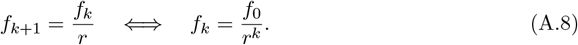

then

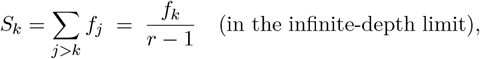

hence the condition *f*_*k*_ *>* 2*S*_*k*_ is equivalent to *r >* 3

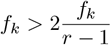

or *r* − 1 *>* 2. For finite depth 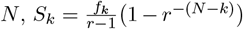, so *r* ≳ 3 suffices in practice with slack set by depth, bandwidths, and modulation indices. More precisely,

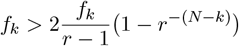

or

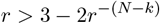

**Figure A.2:**
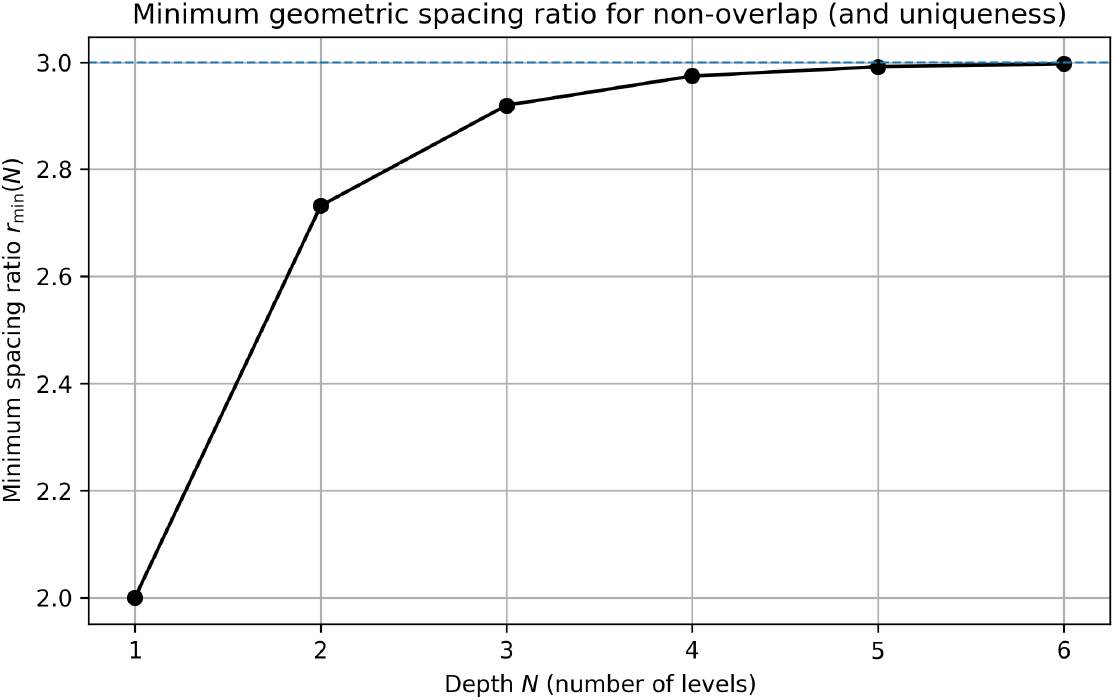
Minimal spacing ratio to avoid overalap as a function of the number of modulation layers. The case *N* = 1 represents the simples, with a carrier and modulation signal (modulation depth is 1).

#### Minimum geometric spacing for finite cascades

For a depth-*N* geometric ladder *f*_*k*+1_ = *f*_*k*_*/r* (*k* = 0, …, *N*), the strongest cluster non-overlap and signed-representation uniqueness condition *f*_*k*_ *>* 2 Σ_*j>k*_ *f*_*j*_ reduces to a single tight inequality at the top level:

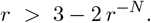

Hence the minimum admissible ratio *r*_min_(*N*) is the unique root *>* 1 of *r* = 3 − 2*r*^−*N*^ . In the infinite-depth limit this yields *r >* 3; for finite depth, *r*_min_(*N*) approaches 3 from below and bound *r*^SI^ (*N*) solving *r* = 2 − *r*^−*N*^, which tends to *r >* 2 as *N* → ∞. is well approximated by *r*_min_(*N*) ≈ 3 − 2*/*3^*N*^ (error *<* 10^−2^ for *N* ≥ 4). Explicitly 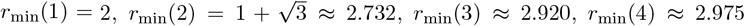, and *r*_min_(5) ≈ 2.992 (see Figure A.2). For comparison, the weaker super-increasing constraint *f*_*k*_ *>*Σ ^*j>k*^ *f*_*j*_ admits the bound 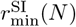 sloving 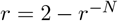 which tends to *r* > 2 as *N* → ∞

#### Remark (Why *f*_*k*_ *> >*Σ ^*j>k*^ *f*_*j*_ is *not* enough)

The weaker “super-increasing” condition *f*_*k*_ *>*Σ ^*j>k*^ *f*_*j*_ guarantees that no *purely slower* combination reaches *f*_*k*_, but it does not exclude overlap of the *clusters* because C_*k*_ extends down to *f*_*k*_−*S*_*k*_ while C_*k*+1_ extends up to *f*_*k*+1_+*S*_*k*+1_ = *S*_*k*_. When *f*_*k*_ ≤ 2*S*_*k*_, these bounds meet or overlap. For example, with *f*_1_ = 6.25, *f*_2_ = 2.5, *f*_3_ = 1 (*r* = 2.5) we have *S*_1_ = 3.5, so C_1_ = [2.75, 9.75] and C_2_ = [1.5, 3.5] overlap on [2.75, 3.5] despite *f*_1_ *> S*_1_.

#### Constant-*Q* and practical benefits

Geometric spacing yields equal steps on a log-frequency axis and approximately constant fractional bandwidth across levels (constant-*Q*), which (i) preserves separability of sideband clusters, and (ii) enables scaleable filter/demodulator design. This mirrors classical constant-*Q* filterbanks and cochlear spacing,^36^ and matches the empirical near-logarithmic arrangement of canonical neural bands.^19, 20^

Log spacing in (A.8) affords constant-*Q* filtering — meaning each band’s 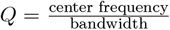 is identical —, so the signal is naturally demodulable in stages: bandpass around *f*_0_ and apply an envelope detector to recover a composite modulation dominated by {*f*_1_, *f*_2_, …}; iterating (envelope of envelope) successively reveals {*f*_1_, *f*_2_, …} at lower and lower centers. The analytic-signal formalism

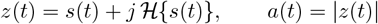

provides a standard implementation of amplitude extraction at each stage.^49^

This hierarchical modulation structure results in nested envelopes that impose amplitude fluctuations at different timescales. As seen in the generated hierarchical modulation plots (see Figure 2.2), a low-frequency modulator introduces a broad envelope variation, a higher-frequency modulator superimposes faster oscillations within that envelope, and so forth. The final signal thus encapsulates a multi-timescale structure where rapid oscillations are embedded within progressively slower amplitude modulations.

#### Bandwidth and modulation index considerations

In a multi-layer design, each modulation layer *k* will effectively use up a certain bandwidth around each spectral line it modulates. If *m*_*k*_ is large (deep modulation), the sidebands carry more power and one might allow a slightly larger bandwidth occupancy for that layer (for example, if the modulating signal is not a pure tone but has its own small bandwidth). However, too large *m*_*k*_ can cause nonlinear distortion (for AM, *m*_*k*_ *>* 1 leads to signal clipping/inversion), which can create additional unwanted spectral lines. Thus, typically one keeps *m*_*k*_ ≤ 1 and often *m*_*k*_ ≪ 1 for higher layers so that those sidebands remain small and confined. The no-overlap condition imposes a trade-off: higher-layer modulations can occupy only a fraction of the spacing left by the layer above.

Natural systems that exhibit hierarchical oscillations appear to enforce the *m <* 1 principle to maintain functional integrity. The brain produces many oscillatory nested rhythms (delta, theta, alpha, beta, gamma, etc.). Maintaining *m <* 1 in these neural modulations preserves **well-separated oscillatory bands**. Each brainwave band retains its characteristic frequency range and identity, which is important if different bands are to carry separable information streams.

In summary, *overmodulation is avoided in both engineered HAM systems and natural oscillatory hierarchies* because it leads to distortion and loss of information. The condition *m <* 1 keeps each modulation layer linear and the overall signal decomposable into distinct frequency components.

### A.2 Power scaling

#### Sideband spectra

We first examine sideband spectra. With equal depths *m*_*i*_ ≡ *m* ∈ (0, 1) and log spacing *f*_*k*+1_ = *f*_*k*_*/r* (*r >* 1), the amplitude of a component involving *k* modulators scales as (*m/*2)^*k*^ . Because *f*_*k*_ ∝ *r*^−*k*^, we have *k* ≈ log_*r*_(*f*_1_*/f*_*k*_). Hence

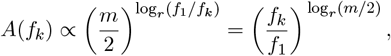

and the power scales as

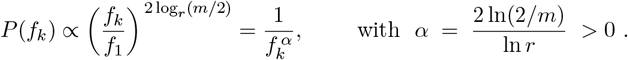

Thus, for fixed *r, α* increases as *m* decreases (weaker modulation ⇒ steeper slope), and for fixed *m, α* decreases as *r* increases (wider spacing ⇒ shallower slope, see Figure 2.4).

#### Full spectrum

Next, we model a random ensemble of logarithmic oscillators interacting via modulation. Consider an ensemble of narrowband oscillators with center frequencies *f* ∈ [*F*_min_, *F*_max_]. Assume (i) *log-uniform* center density *ρ*(*f*) = *C/f* (equal expected number per log-frequency bin, i.e., equal energy per octave), where *C* = [ln(*F*_max_*/F*_min_)]^−1^; (ii) each oscillator contributes the same unit power, shaped by a unit-area spectral kernel *K*_*ε*_(·) that is narrow relative to its center (ideally constant-*Q*, i.e., fractional bandwidth approximately constant across *f*). Equal energy per octave ⇒ pink noise (1*/f*) is a classical result in audio/time–frequency analysis; constant-*Q* filterbanks formalize this *logarithmic* tiling.

##### Lemma 1

**(Log-spaced ensembles yield a 1*/f* backbone).**

Let *S*_0_ denote the ensemble power spectral density (PSD) obtained by summing all oscillators:

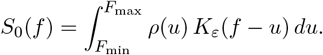

If *K*_*ε*_ is narrow (or constant-*Q*), then, for *f* away from band edges,

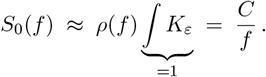

##### Intuition

Convolving a slowly varying density *ρ* with a narrow kernel evaluates *ρ* locally. With *ρ*(*f*) ∝ 1*/f*, equal oscillator mass per octave gives a pink (1*/f*) baseline.

We now assume that the oscillators are coupled randomly as carrier and modulator (second-order pairing). We draw unordered pairs (*f*_*c*_, *f*_*m*_) i.i.d. from *ρ* with the *AM constraint f*_*m*_ *< f*_*c*_. First-order AM generates sidebands at *f*_*c*_ ± *f*_*m*_ with weights *w*_*±*_ (for small modulation depth, *w*_*±*_ ∝ *m/*2).

###### Lemma 2 (Sidebands from a log-uniform population)

Let *f*_*c*_, *f*_*m*_ be i.i.d. on [*F*_min_, *F*_max_] with density *ρ*(*f*) = *C/f*, conditioned on *f*_*c*_ *> f*_*m*_. Define lower/upper sideband frequencies *f*_*L*_ = *f*_*c*_ − *f*_*m*_ and *f*_*U*_ = *f*_*c*_ + *f*_*m*_. Then

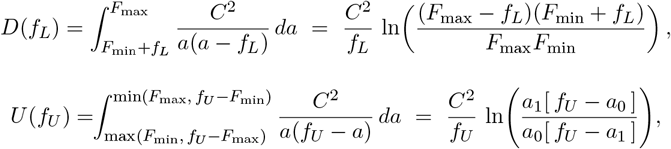

where *a*_0_, *a*_1_ are the integration limits above. For *f* in the interior of [*F*_min_, *F*_max_] both marginals satisfy

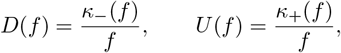

with *κ*_*±*_(*f*) bounded and slowly varying (logarithmic) functions of *f* . As 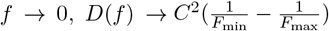 (a constant pedestal), while *U* (*f*) exhibits a high-frequency roll-off near *F*_max_ due to finite support. Thus the *expected* sideband contribution preserves the 1*/f* backbone in the interior, with deviations driven by a low-*f* pedestal (flattening fitted slopes) and a high-*f* truncation (steepening locally).

###### Corollary 1 (Total PSD)

With carrier lines *S*_0_(*f*) ∝ 1*/f* and sidebands *S*_*±*_(*f*) as above, the expected total PSD is

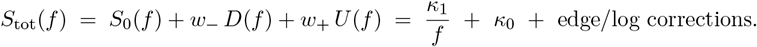

Hence the ensemble remains approximately 1*/f* with an *additive constant κ*_0_ determined by small-difference sidebands and finite-band edges. Larger *κ*_0_ flattens the apparent slope *α* when fitting over restricted ranges.

#### Finite-range and bandwidth effects

(i) *Finite support* [*F*_min_, *F*_max_] produces curvature near edges: high-frequency sum-sidebands are truncated at *F*_max_, and very small differences accumulate near *F*_min_, both tending to *flatten* slopes. (ii) Using a *constant absolute* kernel width (e.g., Gaussian *σ* = 0.5 Hz) is not constant-*Q* and biases the high-frequency region. Adopting *σ* = *ηf* (fractional width) restores scale invariance and improves linearity of log–log fits. These two points explain why, in our simulation, log-spaced centers produce a clear 1*/f* (*α* ≈ 1.0) without sidebands, while adding sidebands slightly reduces the fitted *α* (constant pedestal added), as observed in Fig. 2.3.

#### Does geometric spacing matter for the 1*/f* law?

The derivation of *S*(*f*) ∼ 1*/f* relies only on the *density* of oscillator centers *ρ*(*f*), not on exact geometric ratios. Geometric spacing (*f*_*k*+1_ = *f*_*k*_*/r*) yields *ρ*(*f*) ∝ 1*/f* exactly, but any monotone sequence that satisfies the *super-increasing* condition

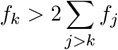

implies that the ratios *r*_*k*_ = *f*_*k*_*/f*_*k*+1_ are all *>* 3, though not necessarily constant. This is because

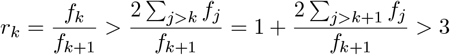

Writing log *f*_*k*+1_ = log *f*_*k*_ − log *r*_*k*_, the sequence of log-frequencies performs a random walk with mean step −⟨log *r*⟩. By the law of large numbers,

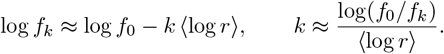

The cumulative count of oscillators above frequency *f* is therefore

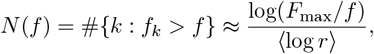

so the differential density is

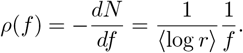

Hence, any super-increasing sequence with bounded ratios and finite ⟨log *r*⟩ yields an approximately log-uniform distribution of oscillator centers. The strict geometric case (*r*_*k*_ ≡ *r*) is a special instance where *ρ*(*f*) = *C/f* exactly. Thus, the weaker non-overlap condition suffices—geometric spacing merely provides a clean analytical form. Thus, geometric spacing is a *sufficient* but not *necessary* condition; the weaker non-overlap constraint already implies an asymptotically log-uniform distribution and therefore the same 1*/f* scaling in expectation — geometric spacing merely provides a clean analytical form.

As we saw, for a *single multiplicative cascade* with geometric spacing *r >* 1 and modulation depth *m*, the toy analysis gives *P* (*f*) ∼ *f*^−*α*^ with *α* = 2 ln(2*/m*)*/*ln *r* (derived earlier). That mechanism ties *α* to (*r, m*) within one chain. By contrast, the *population mixture mechanism* above yields *α* ≈ 1 from the log-spaced *distribution of centers* alone, even without sidebands. Both mechanisms can coexist in neural data; careful parameterization is needed to separate aperiodic from oscillatory peaks.On disentangling aperiodic and periodic structure, see Donoghue et al. (2020).^21^ Other accounts of 1*/f* ^*α*^ (mixtures of timescales, synaptic filtering, E/I balance, etc.) are reviewed in Bédard & Destexhe (2009) and Milotti (2002).^81^

In summary, (1) A bank of log-spaced oscillators with equal power per band produces *S*(*f*) ≈ 1*/f* . (2) Random AM pairings add a constant pedestal; stronger modulation (*m*) or more pairs increase *κ*_0_ and flatten the fitted *α* over finite windows. (3) Using constant-*Q* kernels (or filters) should push the fitted slope in Fig. 1 toward *α* ≈ 1 and increase the fitting range. The assumption *ρ*(*f*) ∝ 1*/f* parallels empirical *logarithmic spacing* of neural bands (delta→gamma)^19^ and is mathematically equivalent to a constant-*Q* tiling (equal spacing on a log-frequency axis).^36^

#### Modulation and demodulation

Let the transmitted field be a narrowband carrier *c*_0_(*t*) around *f*_0_ whose amplitude is multiplicatively structured by slower processes,

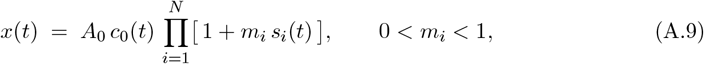

with *s*_*i*_(*t*) narrowband around *f*_*i*_ ≪ *f*_0_. Because (A.9) is a *product*, the *order of modulation* is immaterial: multiplication commutes, hence any staging of the factors yields the same *x*(*t*) (up to higher-order 𝒪(*m*^2^) cross-terms). In network form, convergent inputs *sum then modulate* a faster carrier at a node, but in the small-signal regime, the linear expansion [1 + Σ _*i*_ *m*_*i*_*s*_*i*_(*t*)] reproduces the same product across stages to first order.

To *recover* the nested envelopes from a single wideband signal, order now *does* matter. Write a constant-*Q* demodulator centered at *f* as

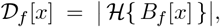

where *B*_*f*_ [·] band-passes the (non-overlapping) cluster around *f* and ℋ{·} is the analytic (Hilbert) transform envelope.^49^ Under the spacing condition *f*_*k*_ *>* 2 Σ _*j>k*_ *f*_*j*_ (equivalently geometric *f*_*k*+1_ = *f*_*k*_*/r* with *r >* 3), clusters do not overlap. Then demodulating the *fastest* band first yields the composite slow envelope,

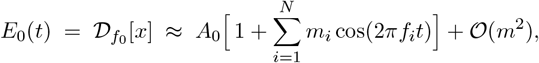

and iterating “fast → slow” peels off deeper layers,

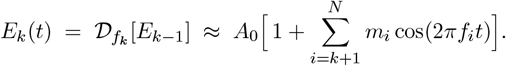

Attempting “slowest-first” on the raw signal is effectively a low-pass, and misses the intended envelope until the faster band that *carries* it has been demodulated. Equivalently, a *parallel* constant-*Q* filterbank can demodulate all layers at once; under non-overlap, these envelopes match the staged fast→slow recovery to first order.

**Terminology: guard-bands, spacing, and non-overlap**

###### Terminology

- **Non-overlap (default meaning)**. Throughout the paper, *“non-overlap”* means **Guard–Band Non–Overlap** (GBNO):

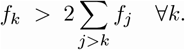 Under *geometric spacing*, i.e., *f*_*k*+1_ = *f*_*k*_*/r*, this is equivalent to *r >* 2–3.
- **Weaker condition**. The **Super–Increasing** (SI) hierarchy

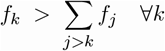

(geometric case *f*_*k*+1_ = *f*_*k*_*/r* ⇒ *r >* 2) prevents slower combinations from *reaching f*_*k*_ but *does not* guarantee that the downward spread of the *k*th sideband cluster cannot overlap the upward spread of the (*k*+1)st. Hence SI is *insufficient* for strict cluster separation and uniqueness of signed combinations.
- **Geometric/log spacing and constant-***Q*. We use *geometric spacing* (*f*_*k*+1_ = *f*_*k*_*/r*) to describe log-uniform centers and a constant-*Q* tiling. When we discuss theoretical guarantees (non-overlap, uniqueness, staged demodulation), we state *r >* 2–3. When we describe empirical bands, we state *r* ≈ 2–3 and note that modest overlap can appear in practice for finite depth and small modulation.

###### Demodulation requirement

- *Sequential demodulation without interference* assumes GBNO (*r >* 2–3 under geometry), so that each carrier’s sideband cluster is disjoint from adjacent clusters.
- When referring to “adequate separation” in simulations or data where depth is finite and modulation is moderate (*m <* 1), we may note that SI (*r >* 2) can suffice operationally, but we reserve the term *“non-overlap” strictly* for GBNO.

###### Summary of statements

- *Theory:* Cluster non-overlap (GBNO) requires *f*_*k*_ *>* 2 Σ_*j>k*_ *f*_*j*_ ∀*k*, i.e., *f*_*k*+1_ = *f*_*k*_*/r* ⇒ *r >* 2–3.
- *Weaker statement:* The super-increasing hierarchy *f*_*k*_ > _*j>k*_ *f*_*j*_ ∀*k* (geometric: *f*_*k*+1_ = *f*_*k*_*/r* ⇒ *r >* 2) prevents slower combinations from reaching *f*_*k*_ but does not preclude adjacent cluster overlap.
- *Empirical:* Canonical bands are roughly log-spaced with *r* ≈ 2–3, close to but sometimes below the theoretical *r >* 2–3, which explains limited yet tolerable overlap in practice.

**Table A.1:** Generators and mechanisms across canonical bands, with neural mass models. Cortical columns can locally generate *γ* (PING/ING) and parts of *α*; thalamo-cortical loops contribute to *α* and dominate *δ*; septo-hippocampal loops pace *θ*; SOs (*<* 1 Hz) primarily emerge from cortical E–I networks with slow processes and can be coordinated by thalamus. Neural mass/field models (JR; corticothalamic mean-field; canonical microcircuits; cortico-basal ganglia) have been proposed for each band: JR yields ∼ alpha and, under parameter shifts, slower (delta-like) rhythms Grimbert (2006); *<* 1 Hz SOs generally require slow adaptation/STD and/or explicit corticothalamic modules Schellenberger Costa (2016), Jajcay (2022). *Notes*. “Very slow” here refers to slow oscillations (SOs, *<* 1 Hz). Infraslow (≲ 0.1 Hz) fluctuations often reflect neurometabolic/vascular components modulating neural excitability and band-limited envelopes rather than a distinct neuronal rhythm (e.g., Hughes (2011), Drew (2020)).

| Band (Hz) | Putative generators | Mechanistic account (abridged) with links (incl. neural mass implementations) |
| --- | --- | --- |
| **Very slow / Slow** (<1) | Neocortex → thalamus (global) | Cortical Up/Down dynamics from recurrent E–I circuits with slow adaptation (e.g., $I_M$, $\text{Ca}^{2+}$-activated $\text{K}^+$) and synaptic depression give bistability and traveling slow waves; cortex alone can generate SOs, with thalamus coordinating SO–spindle interactions. In vivo/in vitro and modeling support this view Sánchez-Vives (2017), Sánchez-Vives (2020), Dalla Porta (2023), Torao-Angosto (2021). *Neural mass implementations*: corticothalamic NMMs/fields that reproduce SOs and K-complexes Schellenberger Costa (2016), Zhao (2015); SO–spindle coupling in a thalamocortical NMM Jajcay (2022). *JR note*: the standard Jansen–Rit (JR) column model exhibits an $\sim\alpha$ limit cycle via Hopf and can also express lower-frequency (delta-like) cycles under parameter changes Grimbert (2006), Spiegler (2010); however, canonical < 1 Hz SOs (Up/Down) typically require slow variables and/or explicit corticothalamic embedding. |
| **Delta** (1–4) | Thalamocortical loop (TC↔TRN) ± cortex | Hyperpolarized thalamic relay (TC) neurons deinactivate T-type $\text{Ca}^{2+}$ channels; interplay with $I_h$ and TRN GABAergic inhibition yields rebound LTS bursts and cyclical TC–TRN pacing in $\delta$; cortex gates/entrains and links $\delta$ to SOs/spindles. Reviews and mechanisms: Crunelli (2018), Crunelli (2006). *Neural mass implementations*: corticothalamic NMMs/fields producing delta/spindles via relay/TRN dynamics (T-currents, $I_h$) Schellenberger Costa (2016), Roberts (2012); JR can enter a slow (delta-like) limit cycle under specific gains/inputs Grimbert (2006), Spiegler (2010). Cortical $\delta$ can also arise locally in EC via HCN resonance Haam (2023). |
| **Theta** (4–8) | Medial septum → hippocampus; hippocampal–entorhinal circuits; links to frontal cortex | Medial septal GABAergic/cholinergic cells pace hippocampus; rhythmic septal inhibition of interneurons yields disinhibition windows for principal cells; CA3/EC inputs and intrinsic resonance shape phase/frequency. Septal cooling slows $\theta$ and disrupts coordination Petersen (2020); system-level reviews Kocsis (2022). *Neural mass implementations*: septo-hippocampal NMMs capturing $\theta$ modulation and phase precession Cona (2015); recent septo-hippocampal NMM with ACh-θ interactions Pirazzini (2025). (Many hippocampal $\theta$ models are spiking/resonator-based rather than NMMs.) |
| **Alpha** (8–12) | Cortex (supragranular) ↔ thalamus (pulvinar/visual nuclei) | Human laminar/iEEG show superficial cortical generators that lead thalamic alpha and propagate as traveling waves Halgren (2019); in mice, visual thalamus is necessary for an alpha-like quiet-wake rhythm and gates cortical activity Nestvogel & McCormick (2022); pulvinar–cortex feedback can generate/gate alpha Cortés (2021). *Neural mass implementations*: cortical JR column (alpha via Hopf) Grimbert (2006); corticothalamic NMMs/fields producing alpha and split-alpha David&Friston (2003), Becker (2015), Roberts (2012). |
| **Beta** (13–30) | Sensorimotor cortex; basal ganglia–thalamocortical loops (STN–GPe) | In cortex, structured E–I connectivity and long-range feedback support transient $\beta$ bursts/traveling $\beta$ waves Kang (2023); in BG, reciprocal STN–GPe forms an E–I resonant loop generating $\sim 20$ Hz; cortico–BG feedback synchronizes and can pathologically amplify beta in PD Pavlidis (2015); layer-V cortical $\beta_2$ (20–30 Hz) in vitro depends partly on gap junction coupling Roopun (2006). *Neural mass implementations*: cortical E–I NMMs reproducing beta waves/bursts Kang (2023); cortico–BG NMMs and mean-field models for pathological beta and DBS effects Pavlidis (2015), Nevado-Holgado (2014). |
| **Gamma** (30–100+) | Local cortical/hippocampal microcircuits (columns) | PING/ING mechanisms: fast E–I loops with $\text{PV}^+$ basket cells and $\text{GABA}_A$ decay set $\gamma$ period; drive and delays tune slow/fast $\gamma$; typically local and nested by $\theta/\alpha$. Mechanistic reviews: Buzsáki & Wang (2012), Craig & McBain (2024). *Neural mass implementations*: canonical microcircuit NMM/DCM families produce $\gamma$ via superficial E–I loops and fast synaptic kinetics Moran (2013), David&Friston (2003), Sánchez-Todo (2023).<sup>33,34,87,88</sup> |

## Footnotes

1 The term Amateur radio (also known as “ham radio”) refers to the use of designated radio frequency bands for non-commercial purposes: self-training, experimentation, recreation, contesting, and emergency communication. The term “ham” apparently comes from land-wire telegraphy (before wireless), where “ham” was used pejoratively to mean a poor or inept operator (e.g., “ham-fisted”). Then, when amateur wireless (radio) got underway in the early 1900s, professional radiotelegraph operators applied the term “hams” to the amateur enthusiasts, deriving from that earlier usage (Wikipedia). For example, one 1909 citation shows the phrase: “I think he is a ham.”

